# Extracellular vesicles synchronize cellular phenotypes of differentiating cells

**DOI:** 10.1101/2021.03.30.437638

**Authors:** Tomohiro Minakawa, Tetsuya Matoba, Jun K. Yamashita

## Abstract

During embryonic development, cells differentiate in a coordinated manner, aligning their fate decisions and differentiation stages with those of surrounding cells. However, little is known about the mechanisms that regulate this synchrony. Here we show that cells in close proximity synchronize their differentiation stages and cellular phenotypes with each other via extracellular vesicle (EV)-mediated cellular communication. We previously established a mouse embryonic stem cell (ESC) line harboring an inducible constitutively active protein kinase A (CA-PKA) gene and found that the ESCs rapidly differentiated into mesoderm after PKA activation. In the present study, we performed a co-culture of control ESCs and PKA-ESCs, finding that both ESCs rapidly differentiated in synchrony even when PKA was activated only in PKA-ESCs, a phenomenon we named “Phenotypic Synchrony of Cells (PSyC)”. We further demonstrated PSyC was mediated by EVs containing miR-132. PKA-ESC-derived EVs and miR-132-containing artificial nano-vesicles similarly enhanced mesoderm and cardiomyocyte differentiation in ESCs and ex vivo embryos, respectively. PSyC is a new form of cell-cell communication mediated by EV regulation of neighboring cells and could be broadly involved in tissue development and homeostasis.

## Introduction

Embryonic development is an extremely fine-tuned and orchestrated process. Individual cells must differentiate in a coordinated manner to form normal tissues. Numerous studies have investigated signaling pathways and gene regulation mechanisms during development [1]. However, models based on the diffusion of morphogens is not sufficient to explain the synchronized cell differentiation. Limited experimental systems have hampered detailed studies on the synchronization that leads to distinct cellular subsets during development.

Embryonic stem cells (ESCs) are derived from the inner cell mass (ICM) of early blastocysts and can differentiate from the pluripotent state into all three germ layers in vitro, making them one of the most powerful tools for studying the cell differentiation process [2–4]. We previously generated a mouse ESC line, PKA-ESCs, in which protein kinase A (PKA) is constitutively activated by the depletion of doxycycline (Dox-OFF) (Figure 1A) [5]. PKA activation with Dox depletion (Dox-) during ESC differentiation in two-dimensional (2D) culture accelerates the appearance of the three germ layers, especially mesoderm cells through a signal-epigenetic linkage [6]. This cell line system is amenable to manipulating the cell differentiation speed only in PKA-ESCs independent of other cell lines; that is, different differentiation speeds can be intentionally generated between activated PKA-ESCs and surrounding ESCs.

**Figure 1.**
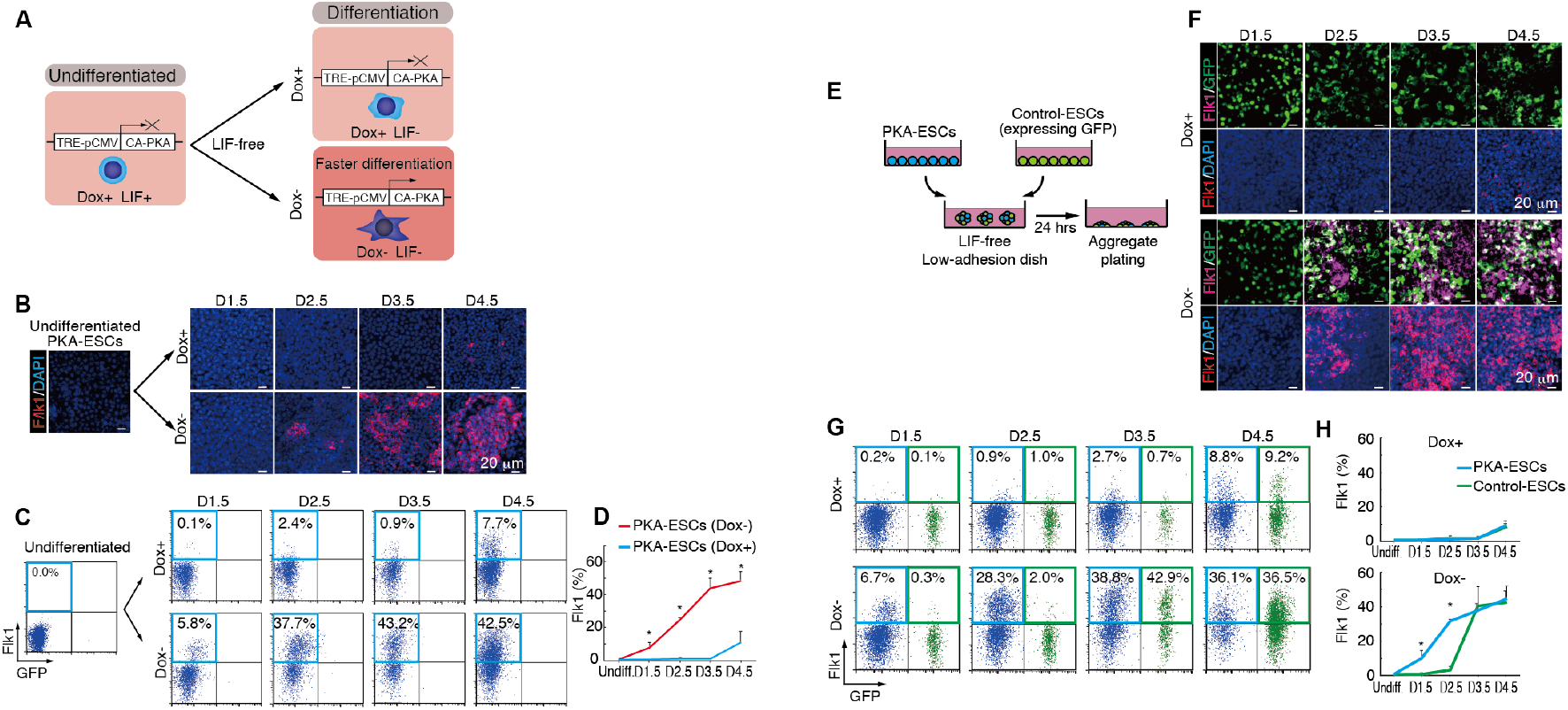
Mouse ESCs synchronize their timing of differentiation with each other. (A) Experimental system of PKA activation. PKA-ESCs express a constitutively active form of PKA (CA-PKA) via a doxycycline-regulated expression system (Dox-Off). To maintain the undifferentiated state, PKA-ESCs were cultured in the presence of LIF and Dox. LIF is excluded for the induction of differentiation. PKA-ESCs differentiate faster in Dox- than in Dox+. (B) Immunostaining for Flk1 during the differentiation of PKA-ESCs. (C) FACS analysis for Flk1 appearance on PKA-ESCs. (D) Percentage of PKA-ESCs expressing Flk1 by FACS analysis. Data were analyzed using the Mann-Whitney U test. Under Dox- conditions, PKA-ESCs showed earlier and significantly more Flk1^+^ cells. *P<0.05 relative to PKA-ESCs (Dox+). (E) Schematic diagram of the chimeric aggregate co-culture differentiation system. PKA-ESCs and Control-ESCs were seeded in low adhesion dishes at a 3:1 ratio to create chimeric aggregates, and differentiation induction was initiated by the depletion of LIF. 24 h later, aggregates were replated on normal plates and attached around 12 h later. (F) Immunostaining for Flk1 during the differentiation of PKA-ESC and Control-ESC chimeric aggregates. The number of Flk1^+^ Control-ESCs in chimeric aggregates (white) increased in Dox- condition. (G) FACS analysis of the Flk1 appearance on chimeric aggregates. Under Dox- condition, Flk1^+^ cells in the PKA-ESC population (blue) appear earlier, but by D3.5, the percentage of Flk1^+^ cells in the Control-ESC population (green) is the same as in the PKA-ESC population. (H) Percentage of chimeric aggregates expressing Flk1 during differentiation. Data represent means ± S.D. Data were analyzed using the Mann-Whitney U test. *P<0.05 relative to Control-ESCs.

In the present study, we performed chimeric aggregate cultures of PKA-ESCs and a control ESC line expressing GFP (Control-ESCs). This experimental system enabled us to intentionally generate different differentiation speeds between two ESC populations in a 3D condition that resembles embryonic development, because the differentiation is specifically accelerated only in PKA-ESCs by Dox depletion. With the use of this unique system, we succeeded in revealing a new phenomenon in which slower differentiating cells synchronize their phenotype with that of faster differentiating cells to reach the same differentiation stage. We name this phenomenon, “<u>P</u>henotypic <u>S</u>ynchrony of <u>C</u>ells (PSyC). We also investigated the molecular machinery that regulates PSyC and identified extracellular vesicles (EVs) and their micro-RNA (miR) content as responsible. PSyC in chimeric cellular aggregates was abrogated by treatment with EV inhibitors. EVs from activated PKA-ESCs promoted the mesodermal differentiation of Control-ESCs and promoted the mesodermal differentiation and even cardiomyocyte differentiation of mouse embryos in ex vivo culture. In addition, among the molecules specifically increased in EVs from PKA-activated ESCs, we identified miR-132 as a functional molecule that can reconstitute the PKA activation and enhanced differentiation of mesoderm lineages in EV-recipient cells. Moreover, a similar cardiomyocyte induction was reproduced by adding artificial nano-particles containing miR-132 to ex vivo mouse embryo culture.

Cell-cell communication via EVs in various tissues has been reported [7]. PSyC offers another mode of this communication and a phenotypic regulation through EVs mainly between cells in close proximity. Our findings provide new understandings about intercellular communication and possible novel strategies towards regenerative medicine.

## Materials and methods

### ESC Lines

PKA-ESC, a mouse ESC line carrying a tetracycline-regulatable constitutively active form of the PKA (CA-PKA) gene, was generated as described previously [5].

EStTA5-4, a mouse ESC line expressing the tetracycline-transactivator protein and containing the puromycin resistance gene [8], was a kind gift from Dr. Takumi Era (Kumamoto University).

Control-ESC (GFP-ESC), a mouse ESC line carrying a ubiquitous promoter-driven EGFP gene, was generated by the introduction of a plasmid carrying the EGFP gene under the control of the CAG promoter to EStTA5-4 cells using the Amaxa Mouse ES cells Nucleofector Kit (VPH-1001, Lonza).

To generate seven mouse ESC lines carrying tetracycline-regulatable miRNAs and GFP, piggyBac vectors carrying primary miRNA genes under the tetracycline response element (TRE) promoter and a transposon vector were introduced using lipofectamine LTX Reagent (15338500, Thermo Fisher Science).

MiR-132 knockout (KO) PKA-ESC line was generated by replacement of genome region of miR-132 with single-strand oligodeoxynucleaotides (ssODNs) including minimal promoter and HiBit. The ssODNs (ordered to Integrated DNA technologies Coralville, USA) and pX330 plasmids expressing Cas9 and a sgRNA targeting the miR-132 locus were introduced into PKA-ESCs using lipofectamine LTX Reagent. Cells were selected by detection of HiBit protein using Nano Glo HiBiT Lytic Detection System (N3030, Promega). pX330 was a kind gift from Dr. Feng Zhang (Massachusetts Institute of Technology).

### Plasmid construction

Primary miR genes were amplified by polymerase chain reaction (PCR) from the genomic DNA of mouse ESCs, and the multi miRs sequence was synthesized by using GENEWIZ. These sequences were cloned into pENTR plasmid vector followed by recombination with piggyBac vector [9] using Gateway LR Clonase II Enzyme Mix (11791-020, Thermo Fisher Scientific). The piggyBac vector and transposon vector were kind gifts from Dr. Knut Woltjen (CiRA, Kyoto University).

### Cell culture and differentiation

ESC lines were maintained in Glasgow Minimum Essential Medium (GMEM; 11710-035, Thermo Fisher Scientific) containing 10% KnockOut Serum Replacement (KSR; 10828-028, Thermo Fisher Scientific), 1% fetal bovine serum (FBS; SAFC Biosciences, USA), 2,000 units/mL leukemia inhibitory factor (LIF; Merck Millipore), 1 mM sodium pyruvate (Sigma-Aldrich), 1% MEM Non-Essential Amino Acids (NEAA; 11140-050, Thermo Fisher Scientific), 50 U/mL penicillin (Meiji Seika Pharma, Japan), 50 μg/mL streptomycin (Meiji Seika Pharma, Japan) and 0.1 mM 2-mercaptoethanol (2-ME; 21985-023, Thermo Fisher Scientific). For the maintenance of PKA-ESCs, 1 μg/mL doxycycline (Dox) was added.

Differentiation was induced in MEM alpha (11900-024, Thermo Fisher Scientific) containing 10% FBS (10437-028, Thermo Fisher Scientific), 2.2 μg/mL NaHCO_3_ (09655-25, Nacalai Tesque, Japan), 50 U/mL penicillin, 50 μg/mL streptomycin and 0.1 mM 2-ME.

To inhibit the secretion of EVs, aggregates were treated with manumycin A (BVT-0091, BioViotica) or GW4869 (13127, Cayman Chemical).

### Aggregate formation

ESC lines were seeded on a low adhesion dish, PrimeSurface Dish 35 mm (MS-9035X, Sumitomo Bakelite, Japan) or low-cell-adhesion 96-well plates with U-bottomed conical wells (9,000 cells per well, 100 mL; MS-9096U, Sumilon PrimeSurface plate, Sumitomo Bakelite) using differentiation medium. After 24 h, floating aggregates were collected and plated on multiwell plates (Falcon). To make chimeric cell aggregates, ESC lines were mixed in ratios as indicated.

### FACS analysis

Cultured cells were harvested at day (D)1.5, D2.5, D3.5 and D4.5 during the differentiation, stained with combinations of allophycocyanin (APC)-conjugated anti-Flk1 mAb (AVAS12) [10] and phycoerythrin (PE)-conjugated anti-platelet-derived growth factor receptor α (PDGFRα) mAb (12-1401-81, eBioscience) and then subjected to analysis with a FACS Aria (Becton Dickinson) or CytoFLEX S (Beckman Coulter).

### Immunohistochemistry

Cultured cells were fixed in ice-cold 4% paraformaldehyde solution for 1 h, blocked in 2% skim milk (232100, Becton Dickinson) for 24 h at 4°C and incubated with primary antibodies for 24 h at 4°C.

The following primary antibodies were used: Flk1 (AVAS12, prepared in house, 1:500,000), PDGFRα (3174S, Cell Signaling Technology, 1:500), Oct4 (sc-5279, Santa Cruz Biotechnology, 1:200) and Nanog (RCAB002P, ReproCELL, Japan, 1:300).

After the primary antibody incubation, the cells were washed with phosphate buffered saline (PBS) three times and incubated with secondary antibodies conjugated to Alexa Fluor (Thermo Fisher Scientific, 1:500) for 24 h at 4°C. Nuclei were visualized with DAPI (4,6 diamidino-2-phenylindole; D3571, Thermo Fisher Scientific).

### RNA-seq for miRNAs

For bulk RNA sequencing, small RNA was extracted using a miRCURY RNA Isolation Kit (300110, EXIQON). We prepared sequencing libraries using the TruSeq Small RNA Library Prep Kit (illumina). The libraries were sequenced in 50 cycle Single-Read mode of HiSeq2500. All sequence reads were extracted in FASTQ format using BCL2FASTQ Conversion Software 1.8.4 in the CASAVA 1.8.2 pipeline. The sequence reads were mapped to mm10 reference genes using MirDeep and normalized and quantified using RPKMforGenes, downloaded on 10 December 2012. Gene expression levels were represented by log2(RPKM+1).

### Western blotting

Isolated EVs were lysed by 2x sample buffer solution without 2-ME (Nacalai Tesque, Japan) and incubated at 70°C for 5 min. The samples were run on sodium dodecyl sulfate-polyacrylamide gel electrophoresis (SDS-PAGE) with the use of a gradient gel (E-T1020L, Atto, Japan), followed by electrophoretic transfer onto nitrocellulose membranes. After the blots, the membranes were incubated for 1 h in the blocking agent Blocking One (03953-95, Nacalai Tesque, Japan), then for 24 h with primary antibodies at 4°C. Anti-mouse or -rat IgG antibodies conjugated with horseradish peroxidase (HRP) were used as the secondary antibodies (1:50,000). The Can Get Signal Immunoreaction Enhancer Solution Kit (NKB-101, Toyobo, Japan) was used for the signal enhancement. Immunoreactivity was detected by using an enhanced chemiluminescence kit, Immobilon Western (WBKLS0500, Merck Millipore). Signal intensities were calculated by using ImageQuant TL Software (GE Healthcare Life Sciences).

### Capillary western blot (WES)

Cell lysates were processed with NE-PER Nuclear and Cytoplasmic Extraction Kit (78835, Thermo Fisher Scientific) to separate the cytoplasmic and nuclear fractions. phosphorylated cyclic AMP response element-binding protein (pCREB), Spry1 and Rasa1 expression levels were measured by using a WES device (Protein Simple).

### ELISA (Enzyme-Linked ImmunoSorbent Assay)

Total CREB and pCREB were quantified using ELISA (85-86153-11, Instant One ELISA CREB1, Thermo Fisher Scientific) in accordance with the manufacturer’s instructions.

### RNA Isolation, RT-PCR (reverse transcription PCR)

Total RNA was isolated from cultured cells using RNeasy (QIAGEN). cDNA was synthesized using SuperScript III (11752-050, Thermo Fisher Scientific) following the manufacturer’s instructions. Quantitative RT-PCR was performed using Power SYBR Green PCR Master Mix (4367659, Thermo Fisher Scientific) and the StepOnePlus system (Thermo Fisher Scientific). Values for each gene were normalized to the relative quantity of GAPDH mRNA in each sample.

### EV isolation

1.0 ×10^5^ PKA-ESCs were seeded in 10 cm dishes with differentiation medium. After 2 days, the medium was changed to a fresh one. After a further 1.5 to 2.5 days of culture, the conditioned medium was collected and centrifuged at 2,000 g for 20 min at room temperature. The supernatant was collected and centrifuged at 12,000 g for 30 min at room temperature. To remove cellular debris and a clump of extracellular matrix, the supernatant was filtered through a 0.2 μm pore filter (Sartorius, Germany) Then the supernatant was ultra-centrifuged at 100,000 g for 60 min at 4°C. EVs were isolated as pellets.

For the in vitro EV-treatment experiments, approximately 1.0 ×10^10^ EVs were added to the cells based on qNano (IZON, UK) analysis. For the ex vivo EV treatment experiments, approximately 1.0 ×10^10^ EVs were added in four aliquots.

### EV Tracking Analysis

EVs were resuspended in PBS, and their size and numbers were analyzed using qNano.

### Transmission electron microscopy analysis

EVs were suspended in PBS (30 μg/mL), dropped (20 μL/drop) on the membrane side of a carbon-stabilized collodion-coated grid (400 mesh; Nisshin-EM, Japan) and left for 10 min at room temperature. The solution was removed with filtered paper and rinsed with distilled water. 1% solution of uranyl acetate dissolved in distilled water was applied to the grid, which was then left to sit for 1 min at room temperature. Then the reagent was removed with filtered paper and dried. The EVs were imaged with a transmission electron microscope (TEM; H-7650, Hitachi High-Technologies, Japan).

### Mice

Pregnant ICR mice on day 3.5 or day 6.5 were purchased from Japan SLC, Inc.

### Embryo ex vivo culture

E6.5 embryos were isolated by dissecting the uterus and decidua in Dulbecco’s Modified Eagle’s Medium (DMEM)/F12 (1:1)(11039-012, Thermo Fisher Scientific) with 5% FBS (SAFC Biosciences, USA). Isolated embryos were cultured on a 12 mm Transwell with a 0.4 μm pore (3460, Corning)(for low-adhesive) with 500 μL DMEM (21063-029, Thermo Fisher Scientific) containing 50% rat serum, 0.1 mM NEAA, 1 mM sodium pyruvate and 0.5 mM 2-ME. After 2 days of culture with or without EVs, the embryos were dissociated with 0.25% trypsin/ethylenediaminetetraacetic acid (EDTA) for 15-25 min at 37°C. The cells were stained with PE-conjugated anti-PDGFRα mAb (12-1401-81, eBioscience) and then subjected to analysis using a FACS Aria (Becton Dickinson).

E3.5 embryos were recovered by flushing uteri with KSOM medium (prepared in house). Embryos were collected into drops of M2 medium (M7167, Sigma-Aldrich). Zona pellucida were removed by brief exposure to acidic Tyrode’s solution drops (T1788, Sigma-Aldrich). M2 medium drop and Tyrode’s solution drops were covered with mineral oil (HiGROW OIL, Fuso Pharmaceutical Industries, Japan) and pre-equilibrated at 37°C with 5% CO_2_. Zona-freed blastocysts were washed with in vitro culture medium (IVC1) (Advanced DMEM/F12 (12634-010, Thermo Fisher Scientific) containing 20% FBS and supplemented with 2 mM L-glutamine (25030-081, Thermo Fisher Scientific), penicillin (25 units/mL)/streptomycin (25 μg/mL), 1x Insulin-Transferrin-Selenium-Ethanolamine (ITS-X) (51500-056, Thermo Fisher Scientific), 8 nM β-estradiol (E8875, Sigma-Aldrich), 200 ng/mL progesterone (160-24511, Wako, Japan) and 25 μM N-acetyl-L-cysteine (A7250, Sigma-Aldrich)) to remove the mineral oil and M2 medium and seeded on a μ-slide 8 well (80826, Ibidi, Germany) filled with pre-equilibrated IVC1 medium as previously described [11]. Two or three days after the embryos were attached on the bottom of the plates, the medium was exchanged to chemically defined IVC2 medium (Advanced DMEM/F12 containing 30% KSR and supplemented with 2 mM L-glutamine, penicillin (25 units/mL)/streptomycin (25 μg/mL), 1x ITS-X, 8 nM β-estradiol, 200 ng/mL progesterone and 25 μM N-acetyl-L-cysteine). The embryo culture was performed at 37°C in 5% CO_2_. The embryos were treated with EVs or polymer poly (DL-lactide-co-glycolide) (PLGA) nanoparticles every two days between day 0 and day 8. All animal experiments were performed in accordance with the guidelines for Animal Experiments of Kyoto University, which conform to the Guide for the Care and Use of Laboratory Animals in Japan.

### Preparation of PLGA nanoparticles

Biotinized PLGA with an average molecular weight of 20,000 Da and a copolymer ratio of lactide to glycolide of 75:25 (Nanosoft Polymers, NC, USA) was used as the matrix for the nanoparticles, and polyvinylalcohol (PVA-403, Kuraray, Japan) was used as the dispersing agent. PLGA nanoparticles incorporating a mirVana miRNA mimic hsa-mir 132-3p (4464066, Thermo Fisher Science) were prepared using an emulsion solvent diffusion method in RNAase-free water, as previously described [12]. The sequence of human miR-132-3p and mouse miR-132-3p are identical. miR-132-PLGA contained 11.6% (w/w) miR-132. A sample of the nanoparticle suspension in distilled water was used to determine the particle size. The median diameter of miR-132-PLGA based on dynamic light scattering was 262 nm.

### Statistical analysis

The significance of differences was evaluated using the Mann–Whitney U test for comparisons between two groups or by the Kruskal–Wallis test, followed by the Steel– Dwass test for multiple comparisons. *P* > 0.05 was considered not significant.

## RESULTS

### Mouse ESCs synchronize their stage of differentiation

In our previous differentiation condition (differentiation medium with serum) in 2D culture, mouse ESCs gave rise to Flk1^+^ mesoderm cells at around 4 days of differentiation (D4) [6,13]. In the present study, first we confirmed the differentiation speed of mouse ESCs in cell aggregates (Figure S1A). Similar to our recent report [6], activated PKA-ESCs (Dox-) in cell aggregates differentiated much more rapidly to Flk1^+^ cells than did non-activated PKA-ESCs (Dox+) (Figure 1B). Quantitatively, we found fewer than 1% of total non-activated PKA-ESCs were Flk1^+^ at D3.5, but more than 40% of activated PKA-ESCs were (Figures 1C and 1D). The appearance of a mesoderm population expressing another mesoderm marker, PDGFRα^+^, was also enhanced upon PKA activation (Figures S1B, S1C and S1D). The differentiation speed of Control-ESCs was not affected by Dox exposure (Figures S1E-S1J). In both Dox+ and Dox-, the percentage of Flk1^+^ cells in Control-ESCs was less than 10% at D3.5 (Figure S1E). We then created chimeric cell aggregates of PKA-ESCs and Control-ESCs (Figure 1E). In this co-culture system, PKA activation can generate different differentiation speeds between the two cell lines; that is, the differentiation is expected to be accelerated only in PKA-ESCs after PKA activation with Dox depletion. Surprisingly, however, in the chimeric aggregates, PKA activation in PKA-ESCs induced an earlier appearance of mesoderm cells not only in PKA-ESCs, but also in Control-ESCs at D3.5 at comparable levels (Figures 1F-1H). Immunostaining for Flk1 (Figure 1F) showed that under Dox-condition, an earlier appearance of Flk1^+^ cells (red) was observed not only in GFP^−^ cells (PKA-ESCs) but also in GFP^+^ cells (control ESCs), that is, considerable numbers of GFP^+^Flk1^+^ cells (white in Figure 1F) appeared from D2.5, suggesting faster differentiation of Control-ESCs in this condition. Flow cytometry analysis (Figure 1G and 1H) more clearly demonstrated the earlier and synchronized appearance of Flk1^+^ cells after Dox depletion. In control condition (Dox+), both PKA-ESCs (blue dots) and Control ESCs (green dots) similarly differentiated into Flk1^+^ cells at comparable levels until D4.5 (Figure 1G). In Dox- condition, which activates PKA only in PKA-ESCs, the appearance of Flk1^+^ cells was evoked in PKA-ESCs from D2.5 and reached a much higher level at D3.5-4.5 than in control condition. As if following the earlier differentiation of PKA-ESCs, Flk1^+^ cells in the Control-ESC population started to appear and eventually caught up in percentage with those in the PKA-ESC population by D3.5, even though Dox depletion did not activate PKA in Control-ESCs (Figure 1G). Consistently, the percentage of Flk1^+^ cells in the Control-ESC population lagged behind that in the PKA-ESC population until D3.5, but their differentiation levels were synchronized thereafter (Figure 1H). A similar early and synchronized appearance of PDGFRα^+^ cells in the Control-ESC population was seen in chimeric cell aggregates after PKA activation (Figures S1K-S1M). Together, these results indicate that there exists a novel cellular mechanism that synchronizes the cellular phenotypes of different cell populations (i.e., PSyC).

### PSyC is mediated by EVs

Next, we examined the mechanism of PSyC. We searched for intercellular communication systems that can conduct PSyC, such as humoral factors, gap junctions, and EVs, finding EVs as a potent candidate. EVs include exosomes and microvesicles (also called ectosomes). Exosomes are 50-150 nm-sized EVs that are generated by the inward budding of endosomes [14]. Microvesicles are 100-1000 nm-sized EVs that are generated by the direct budding of plasma membranes [15,16]. Cells transfer several types of biomolecules, including mRNA, miR and proteins, via EVs [7]. We initially examined the loss-of-function of EVs by inhibiting EV secretion (Figure 2A).

**Figure 2.**
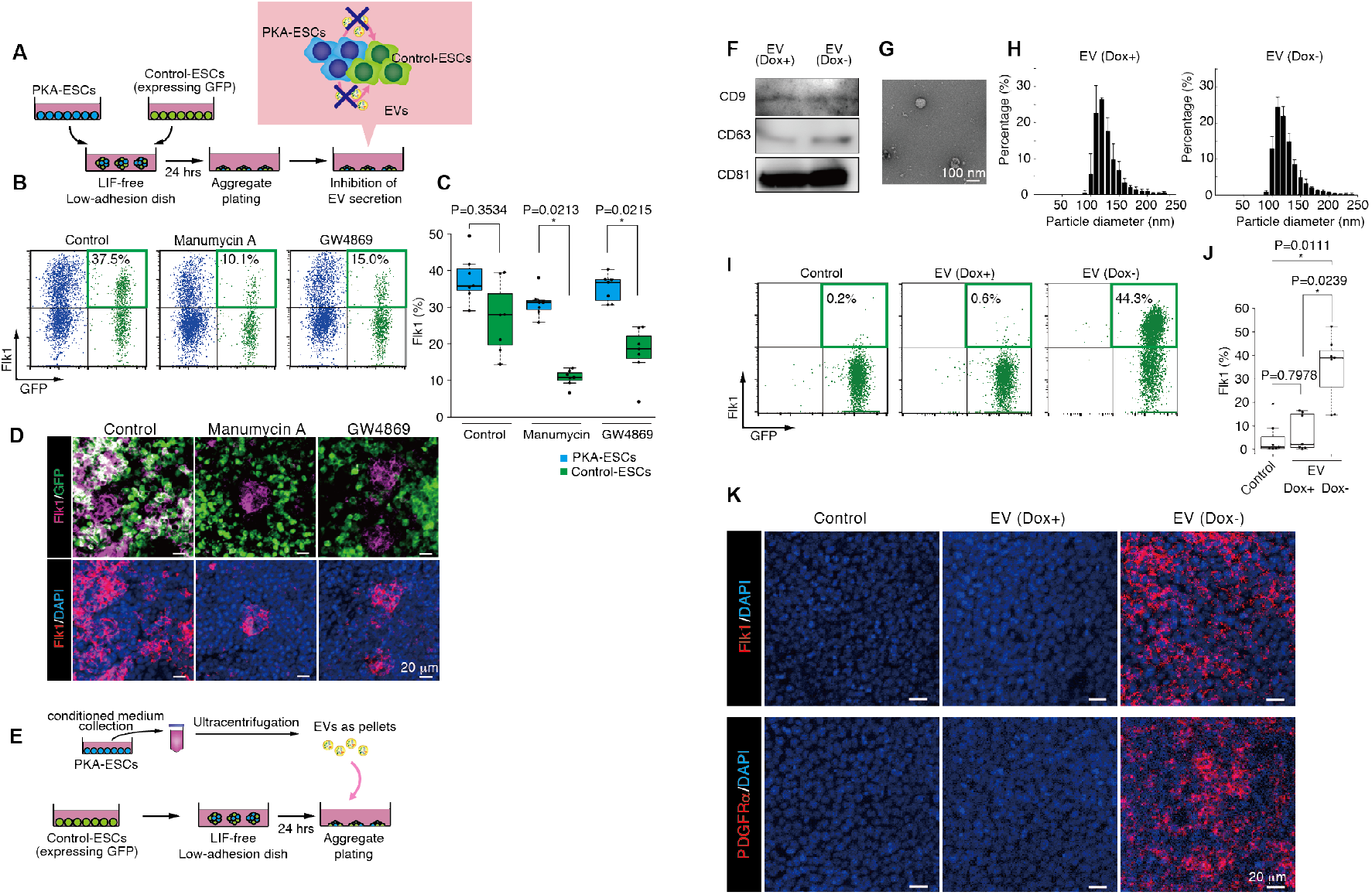
PSyC is mediated by EVs. (A) Schematic diagram of the chimeric aggregate co-culture differentiation system with EV secretion inhibited. PKA-ESCs and Control-ESCs were seeded in low adhesion dishes at a 3:1 ratio to create chimeric aggregates, and differentiation induction was initiated by the depletion of LIF. 24 h later, the aggregates were replated on normal plates and cultured with EV secretion inhibitors. (B) The number of Flk1^+^ cells was reduced with treatment of the EV secretion inhibitors Manumycin A (10 μM) or GW4869 (5 μM). PKA-ESC and Control-ESC chimeric aggregates on D3.5 under Dox- condition were analyzed by FACS. (C) Percentage of PKA-ESCs and Control-ESCs in chimeric aggregates expressing Flk1 under Dox- condition. Data were analyzed using the Kruskal-Wallis test followed by the Steel-Dwass test. (D) Immunostaining for Flk1^+^ Control-ESCs (white) in chimeric aggregates with manumycin A (10 μM), GW4869 (5 μM), or DMSO (control). (E) Schematic diagram of the EV collection and treatment. Control-ESCs aggregates were plated without LIF. EVs were collected from the conditioned medium of PKA-ESCs as pellets and added to Control-ESCs aggregates. (F) Immunoblots of CD9, CD63 and CD81 in the exosomal protein lysate from 10 mL conditioned medium. (G) Transmission electron microscopy (TEM) image of EVs from the conditioned medium. (H) Analysis of the EV size distribution in Dox+ (left) and Dox- (right) conditioned medium. (I) FACS analysis of Flk1^+^ cells in EV-treated Control-ESCs on D3.5. (J) Percentage of Control-ESCs expressing Flk1. Data were analyzed using the Kruskal-Wallis test followed by the Steel-Dwass test. (K) Immunostaining for mesodermal markers (Flk1, PDGFRα) in EV-treated Control-ESCs on D3.5. Data represent means ± S.D.

Manumycin A and GW4869 are two inhibitors of enzyme neutral sphingomyelinase 2 (nSMase2), which regulates exosome secretion. Either manumycin A or GW4869 treatment specifically and significantly decreased the expression of Flk1^+^ cells only in the Control-ESC population, without affecting PKA-ESCs in chimeric aggregates under PKA activation at D3.5 (Figures 2B and 2C). Immunostaining revealed that the expression of Flk1 by Control-ESCs was decreased by the inhibitor treatment (Figure 2D, white). These results suggested the involvement of EVs in PSyC. To analyze the effect of EVs secreted from PKA-ESCs, we next isolated EVs in a conditioned medium of PKA-ESCs under non-active (EV (Dox+)) and active (EV (Dox-)) PKA conditions (Figure 2E). Immunoblotting of the EV markers CD9, CD63 and CD81 (mainly expressed in exosomes and sometimes in microvesicles) confirmed EV isolation (Figure 2F). EVs were observed as vesicles 100-150 nm in diameter by electron microscopy and EV tracking analysis (Figures 2G and 2H). The particle numbers and protein concentrations in EV (Dox+) and EV (Dox-) were not significantly different (Figures S2A and S2B). The addition of EV (Dox-) to pure Control-ESC aggregates induced a much higher percentage of Flk1^+^ and PDGFRα^+^ cells than no EVs or EV (Dox+) (Figures 2I, 2J, S2C and S2D). Consistently, immunostaining showed a dramatic increase in Flk1 and PDGFRα expression after EV (Dox-) treatment (Figure 2K). These results indicate that EVs from activated PKA-ESCs enhance the differentiation of Control-ESCs to mesoderm to synchronize the differentiation levels.

### EV-derived miRs regulate PSyC

Next, we tried to identify molecules within the EVs that induced PSyC. Global miR expression profiles contained in EV (Dox+) and EV (Dox-) were compared by miR-seq (Figure 3A). We found that miR-126, −132, −184, −193a, −212 and −483 were enriched in EV (Dox-). We generated ESC lines overexpressing all 6 miR (multi miR) or each miR together with GFP under a Dox-OFF system and performed cell chimera experiments. The parental mouse ESC line [8] for the miR-overexpressing cells does not carry any miR or GFP genes and was used as the recipient cells. In order to specifically evaluate the effects of miRs transferred from miR-expressing cells (GFP^+^) to recipient cells (GFP^−^), we analyzed mesoderm induction only in recipient cells (Figure 3B). We generated chimeric cell aggregates with recipient cells and each miR-expressing cell line at a 1:3 ratio. Flk1^+^ mesoderm differentiation in the recipient cells (GFP^−^) was significantly promoted only when mixed with the miR-132 or multi miR cell line (Figures 3B and 3C). We further confirmed the role of EV-transferred miR-132 in the mesoderm induction of the recipient cells by generating miR-132 KO PKA-ESCs. EVs were prepared from the supernatants of PKA-ESCs or miR-132 KO PKA-ESCs after PKA activation (Dox-) and added to the differentiating recipient cells. Whereas EVs from PKA-ESCs with Dox- (EV (Dox-)) enhanced the mesodermal differentiation of the recipient cells (similar to Figure 2I), EVs derived from miR-132 KO PKA-ESCs (Dox-) failed to promote mesoderm differentiation (Figures S3A and S3B). These results indicate that miR-132 in EV (Dox-) is a major messenger molecule in the enhancement of mesoderm differentiation in recipient cells and contributes to PSyC.

**Figure 3.**
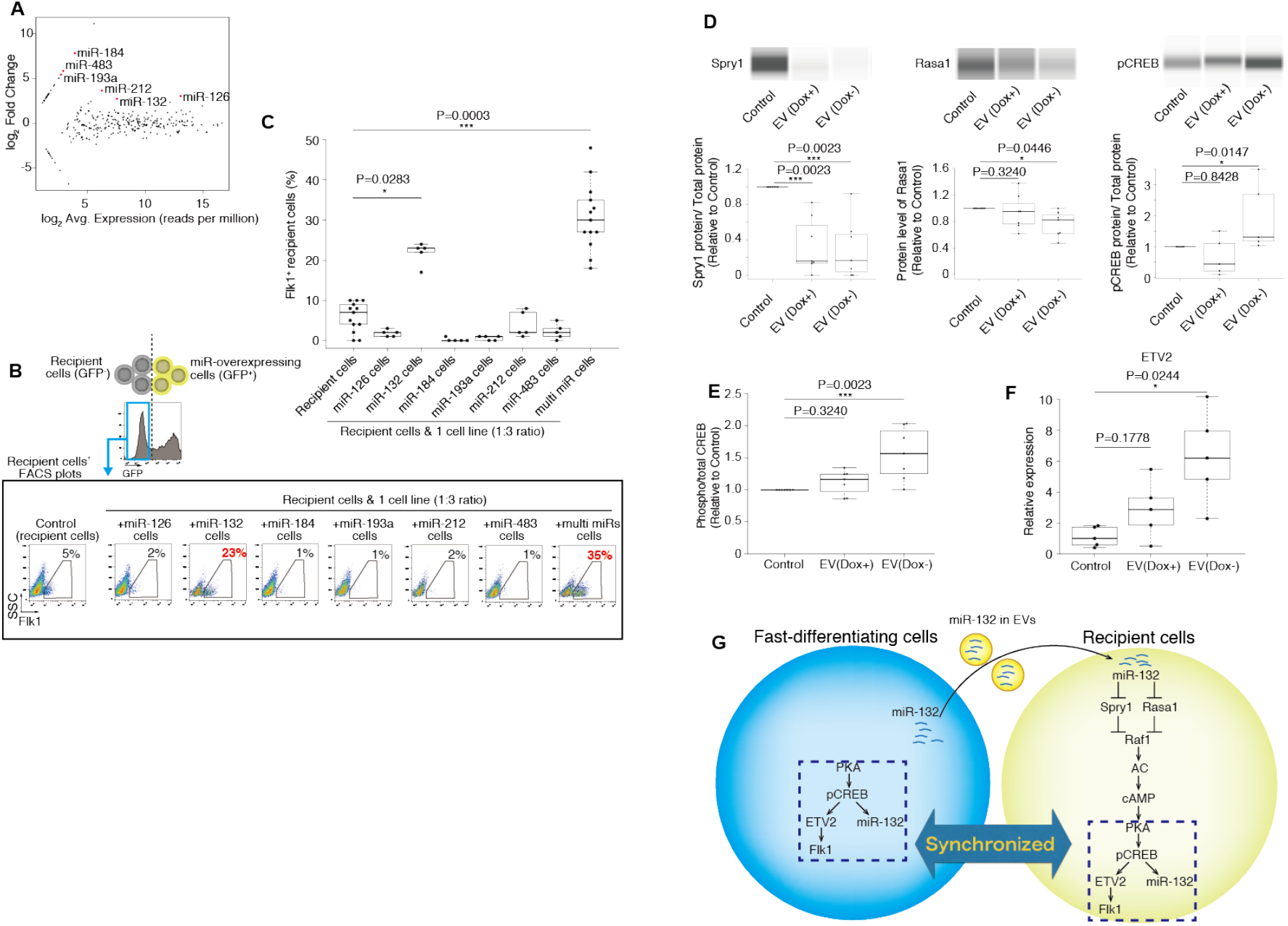
EV-derived microRNAs regulate PSyC. (A) An MA plot summarizing the differentially expressed miRNAs in EVs from activated PKA-ESCs (Dox-) versus EVs from inactivated PKA-ESCs (Dox+). (B) Flk1^+^ recipient cells (EStTA5-4) in chimeric aggregates of recipient cells and miRNA-expressing cells (1:3 ratio). FACS plots showing recipient cells in chimeric aggregates with miRNA-expressing cell lines on D3.5. (C) Percentage of Flk1^+^ recipient cells in aggregates at a 1:3 ratio. Data were analyzed using the Kruskal-Wallis test followed by the Steel-Dwass test. (D) Detection of Spry1, Rasa1 and pCREB in capillary western blots. Proteins in the nuclear fractions were analyzed to measure the expression levels of Spry1 and pCREB and normalized with Lamin A/C. Proteins in the cytoplasmic fractions were analyzed to measure the expression level of Rasa1 and normalized with β-actin. Data were analyzed using the Kruskal-Wallis test followed by the Steel-Dwass test. (E) pCREB and CREB levels were measured by ELISA. Data were analyzed using the Kruskal-Wallis test followed by the Steel-Dwass test. (F) qPCR analysis of ETV2 in Control-ESCs treated with EVs from PKA-ESCs (Dox+ or Dox-) or without EVs. Data were analyzed using the Kruskal-Wallis test followed by the Steel-Dwass test. (G) Schematic of the PSyC mechanism including EV secretion and miR-132 delivery. From fast-differentiating cells, miR-132 is passed through EVs to surrounding recipient cells, where it inhibits Spry1 and Rasa1 to transmit the signal. As a result, the differentiation mechanism is synchronized in the fast-differentiating cells and recipient cells. Data represent means ± S.D.

We further examined the intracellular molecular machinery of the PSyC induction in recipient cells after they received EVs that carried miR-132. Spry1, an antagonist for fibroblast growth factor pathways, and Rasa1, a RasGAP activator, are targets of miR-132 [15,17,18]. When Control-ESCs were treated with EV (Dox-), Spry1 and Rasa1 protein levels were significantly reduced, indicating that miR-132 acts in Control-ESCs through EV-mediated transfer (Figure 3D). miR-132 has been reported to activate Ras/Raf1 signaling by inhibiting Spry1 and Rasa1 [17]. Additionally, Raf1 is reported to phosphorylate CREB via the adenylyl cyclase/cyclic AMP/PKA [19,20] and ERK/RSK pathways [21,22]. Furthermore, pCREB upregulates the expression of ETS variant transcription factor 2 (ETV2) [23] and Flk1 [24] genes, both of which are mesoderm inducers, and of miR-132 [25]. Thus, the reduction of Spry1 and Rasa1 protein expression is postulated to activate PKA signaling through pCREB and result in mesoderm differentiation. Indeed, the levels of CREB phosphorylation (Figures 3D and 3E) and ETV2 mRNA (Figure 3F) in Control-ESCs were increased by the addition of EV (Dox-). Taken together, our results suggest a molecular link in PsyC after PKA activation as follows. First, PKA activation initiated in PKA-ESCs enhances Flk1^+^ mesoderm cell differentiation as well as miR-132 expression through CREB phosphorylation. The activated PKA-ESCs then secrete EVs with high miR-132 content. miR-132 is delivered to the recipient cells via EV-mediated transfer and suppresses Spry1 and Rasa1, which in turn activates the PKA pathway through CREB phosphorylation in Control-ESCs. Thus, PKA activation initiated in the original PKA-ESCs can be reconstituted in Control-ESCs through PKA-ESC-derived EVs to evoke PSyC. Following these findings, we concluded that PSyC is a novel biological mechanism that reconstitutes similar intracellular environments in donor and recipient cells in close proximity via donor-derived EVs (Figure 3G).

### PKA-ESC-derived EVs and miR-132 induce mesoderm and cardiac myocytes in mouse embryos

Finally, we examined whether EVs from differentiating ESCs affect cell fate during embryonic development. First we tried to confirm the effect of EV (Dox-) on mesoderm stage embryo. We collected E6.5 mouse embryos and cultured them ex vivo for 2 days with EV (Dox-) treatment (Figure 4A). We dissociated the embryos and examined PDGFRα^+^ cells in the embryos after 2 days of ex vivo culture by FACS. Significantly more PDGFRα^+^ cells were observed in the embryos when treated with EV (Dox-) compared to the untreated group (Figures 4B and 4C), suggesting that EV (Dox-) can enhance mesoderm differentiation in embryonic development.

**Figure 4.**
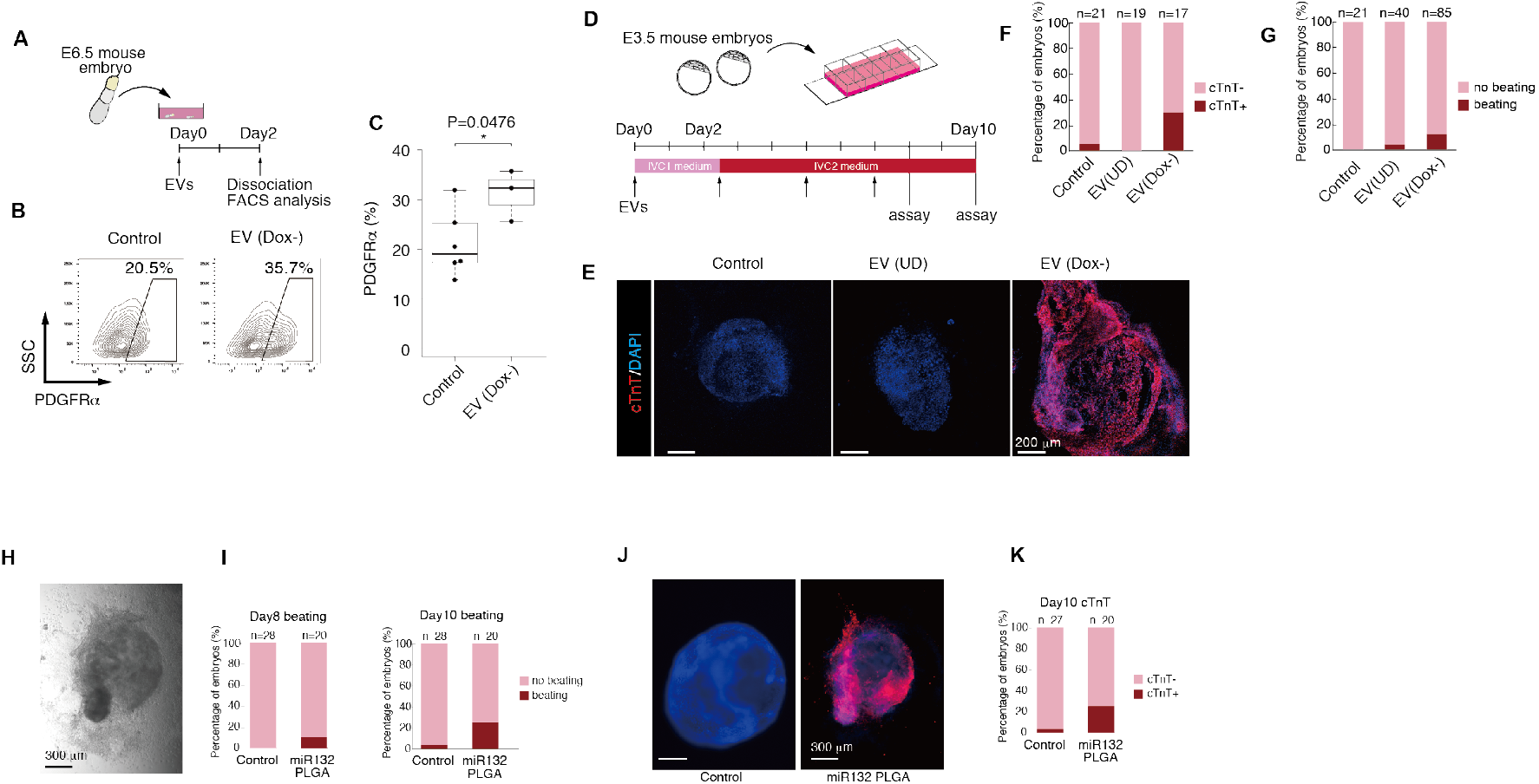
PKA-ESC-derived EVs and miR-132 induce mesoderm and cardiac myocytes in mouse embryos. (A) Schematic diagram of the mouse embryo culture. E6.5 mouse embryos were cultured for 2 days with EV treatment. At day 2, mouse embryos were dissociated and subjected to FACS analysis. (B) FACS analysis of PDGFRα expression on E6.5 embryos cultured for 2 days with EVs isolated from undifferentiated (UD) Control-ESCs or differentiating PKA-ESCs under Dox- condition. (C) Percentage of Control-ESCs expressing Flk1. Data were analyzed using the Mann-Whitney U test. (D) Schematic diagram of the mouse embryo culture. E3.5 mouse embryos isolated from the uterus were cultured on μ slides in IVC1 medium. After attaching the mouse embryos to the slides on days 2-3, the medium was changed to IVC2 medium, and the embryos were cultured until day 8 or 10. EVs were added once every two days. (E) Immunostaining for cTnT on E3.5 mouse embryos cultured for 8 days with EVs isolated from UD Control-ESCs or differentiating PKA-ESCs under Dox- condition. (F and G) Percentage of cTnT^+^ (F) and beating (G) embryos. E3.5 mouse embryos were cultured for 10 days with undifferentiated ESC-derived EVs (EV(UD)) or early differentiating PKA-ESC-derived EVs (EV(Dox-)) (H) A beating mouse embryo cultured for 10 days from E3.5 with miR-132-PLGA. (I) Percentage of beating embryos on day 8 and day 10. (J) Immunostaining for cTnT on E3.5 mouse embryos cultured for 10 days with or without miR-132-PLGA. (K) Percentage of cTnT^+^ embryos on day 10. Data represent means ± S.D.

We further examined the effects of EV (Dox-) on cell fate determination from an earlier stage of development. We applied a recently established ex vivo culture system for the observation of embryos continuously from the preimplantation to postimplantation stages [11]. We collected E3.5 mouse embryos and cultured them ex vivo with EVs (Figure 4D). Surprisingly, culturing the earlier embryos with EV (Dox-) drastically altered the embryo development. The appearance of cardiomyocytes with cardiac troponin-T-positive cells was enhanced in embryos treated with EV (Dox-) (5 of 17 embryos), but never in embryos treated with EVs collected from undifferentiated Control-ESCs (EV (UD)) (0 of 19 embryos) and rarely in untreated embryos (1 of 21 embryos; Figures 4E and 4F). Additionally, whereas just weak beating or twitching was detected in only a few EV (UD)-treated and untreated embryos (2 of 40 embryos and 0 of 21 embryos, respectively), distinct cell clusters with apparent and typical spontaneous beating, sometimes dominating the whole embryo, were observed in EV (Dox-)-treated embryos (11 of 85 embryos; Figure 4G and Supplementary Movie 1). Thus, EV (Dox-) has the potential to alter cell fate to mesoderm derivatives including cardiomyocytes in developing embryos.

We further confirmed whether miR-132 similarly works on cell fate determination. For this purpose, we employed an artificial polymeric nanoparticle that is formulated from biodegradable polymer poly (DL-lactide-co-glycolide) (PLGA) and can entrap various molecules including nucleic acids, penetrate cellular membranes, and deliver the encapsulated content into the cellular cytoplasm. We formulated PLGA nanoparticles containing miR-132 (miR-132-PLGA). miR-132-PLGA added to differentiating mouse ESCs successfully upregulated mesoderm marker Flk1 mRNA expression, indicating that miR-132-PLGA can mimic the function of EV (Dox-) (Figure S3C). When miR-132-PLGA was added to E3.5 mouse embryos, similar to that with EV (Dox-) addition, beating cardiomyocytes were observed from day 8 and increased in number until day 10. The percentage of embryos with beating cardiomyocytes was 3% in Control (N = 28) but as high as 25% in the miR-132-PLGA group (N = 20). (Figures 4H and 4I). All embryos with beating cardiomyocytes were positive for cardiac troponin-T (Figures 4J and 4K). Thus, we confirmed that miR-132 shows the same potential as EV (Dox-) with regards to altering cell fate towards mesoderm and cardiomyocytes in vivo.

## DISCUSSION

We showed a novel biological phenomenon in which cellular phenotypes are synchronized during differentiation by EV-mediated cell-cell communication, “Phenotypic Synchrony of Cells (PSyC)”. We identified one potential candidate molecule in the EVs, miR-132, that can reconstitute the molecular link in the recipient cells for PSyC and reproduce the in vivo effects of EVs to induce mesoderm derivatives.

Though various mesoderm-inducing molecules have been reported previously, such as basic fibroblast growth factor (bFGF), bone morphogenetic protein (BMP), and PKA pathway molecules, our global protein analysis with mass spectrometry showed that these molecules were not enriched in EV (Dox-) (data not shown). Instead, miR-132, which has not been previously reported to induce mesoderm, was enriched and could evoke mesoderm induction in EV-recipient cells through the reconstitution of PKA activation. However, considering that EVs contain thousands of biomolecules, such as miRs and proteins, the observed miR-132-mediated effect should represent only part of the EV-mediated PSyC mechanism. Moreover, PSyC may more broadly contribute to cell and tissue development and homeostasis. The PSyC phenomenon we demonstrated here would be just one type of PSyC effects. We speculate that the molecules responsible for PSyC vary with the cell type and microenvironment. For example, it has been reported that Cav1 KO adipocytes receive Cav1 mRNA from surrounding endothelial cells via EVs, allowing the KO cells to produce Cav1 protein [26]. Accordingly, PSyC with different conditions and environments should be investigated.

Recently, EVs were highlighted as a novel cell-cell communication modality that acts through remote effects analogous to hormones [16]. On the contrary, in PSyC, EVs function at short-range, transmitting the cellular state to surrounding cells in cell aggregates. This synchronizing effect was largely reduced when PKA-ESCs and Control-ESCs were separately cultured with a transwell membrane (data not shown), suggesting that the EVs are absorbed by neighboring cells to exert PSyC. EVs are a heterogeneous group of cell-derived membranous structures including exosomes and microvesicles, which originate from the endosomal system or which are shed from the plasma membrane, respectively [27]. Exosomes are generated by endosomal sorting complex required for transport (ESCRT)-dependent and ESCRT-independent mechanisms [27]. It has been reported that the silencing of tumor susceptibility 101 (TSG101), a component of the ESCRT complex, reduces exosome secretion [28,29]. The generation of arrestin domain containing 1 (ARRDC1)-mediated microvesicles also requires TSG101 [30]. Consistently, it has been reported that TSG101 KO inhibits cell proliferation and differentiation, and TSG101 KO mice show embryonic lethality around the implantation stage [31]. In contrast, mice knocked out of nSMase2, which is involved in ESCRT-independent exosome production, show hypoplasia and growth retardation in all tissues, but no fetal lethality [32]. In our study, PSyC was significantly but not completely blocked by nSMase2 inhibitors, suggesting the involvement of ESCRT-independent pathways but also that of other EV generation mechanisms, ESCRT-dependent exosomes and/or microvesicles.

In this study, we report a new intercellular communication mechanism between cells in close proximity. We speculate that PSyC would establish cell subsets to build normal tissues through a synchronization mechanism that closely aligns cell fate determination and differentiation stages. Further, PSyC should be a critical mechanism for regulating tissue development and homeostasis. Tissues and organs may be established and maintained with more active and massive molecular transport through EV exchange among neighboring cells.

## Acknowledgements

We thank Dr. T. Matsunaga and T. Hoshino for technical suggestions and Dr. P. Karagiannis for critical reading of the manuscript. We thank Dr. A. Watanabe, Dr. M. Nakamura and N. Amano for analyzing the RNA-seq data. We also thank Dr. I. Baba, Kozue Nakamura, Kae Nakamura and Y. Kaji for technical assistance. This work was supported by JST CREST Grant Number JPMJCR17H5, Japan.

## Author Contributions

T.Minakawa and J.K.Y. designed the experiments. T.Minakawa performed the experiments including the cell culture, FACS, qRT-PCR, immunostaining, EV collection, EV analysis and mouse embryo culture. T.Matoba prepared the PLGA nanoparticles. T.Minakawa and J.K.Y. wrote the manuscript. J.K.Y. supervised the study.

---

**Figure S1.**
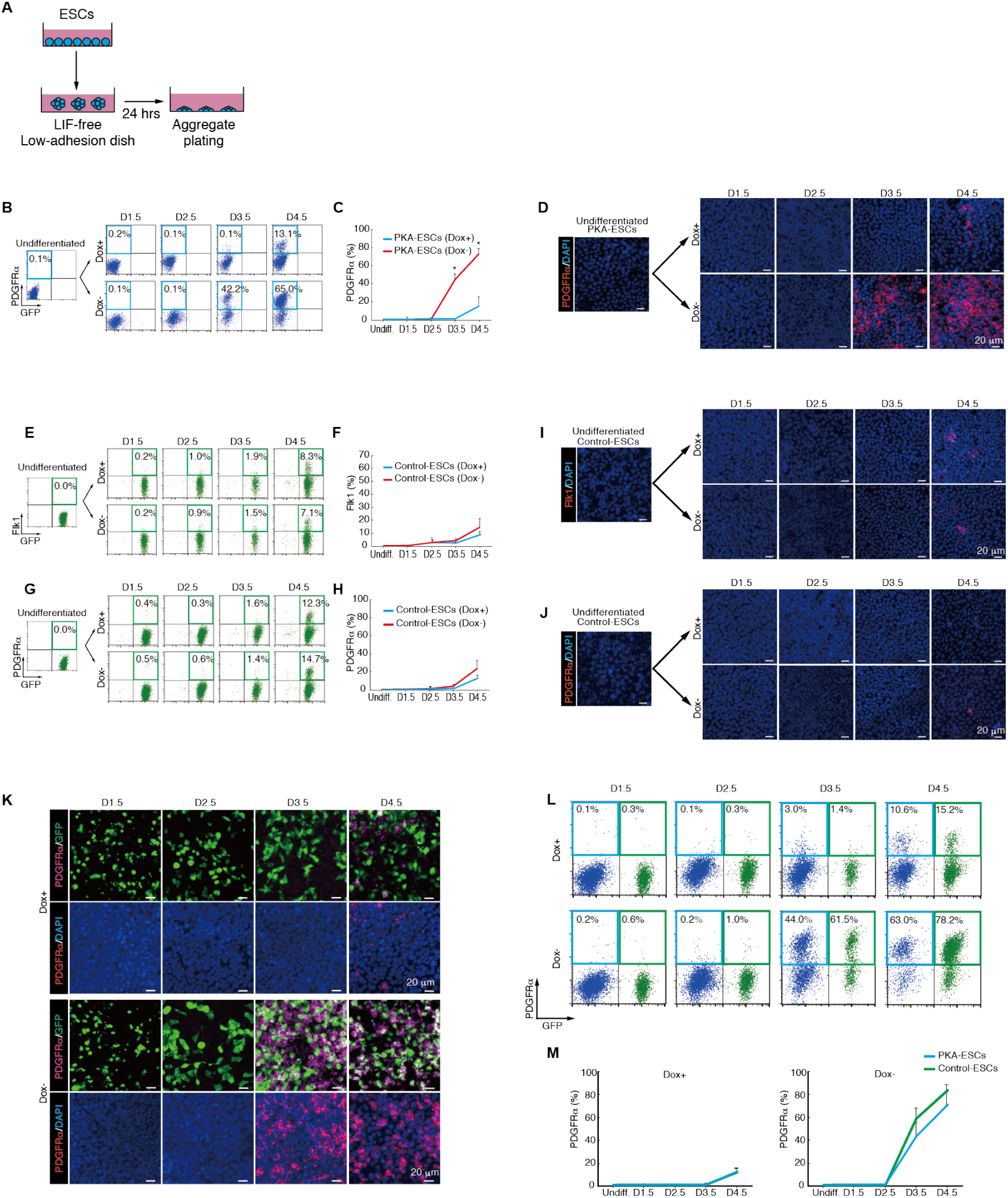
Aggregate culture of PKA-ESCs and Control-ESCs. (A) Schematic diagram of the aggregates culture. Mouse ESCs were seeded in low adhesion dishes to create aggregates, and differentiation induction was initiated by the depletion of LIF. 24 h later, the aggregates were replated on normal plates and attached at around 12 h later. (B) FACS analysis of pure PKA-ESC aggregates for PDGFRα. (C) Percentage of PDGFRα^+^ PKA-ESCs. Data were analyzed using the Mann-Whitney U test. *P<0.05 relative to PKA-ESCs (Dox+). (D) Immunostaining of PDGFRα during the differentiation of pure PKA-ESC aggregates. (E-H) FACS analysis of pure Control-ESC aggregates for Flk1 (E) and PDGFRα (G). Percentage of Flk1^+^ Control-ESCs (F) and PDGFRα^+^ Control-ESCs (H). (I and J) Immunostaining of Flk1 (I) and PDGFRα (J) during the differentiation of pure Control-ESC aggregates. (K) Immunostaining of PDGFRα during the differentiation of chimeric aggregates. The number of PDGFRα^+^ Control-ESCs in chimeric aggregates (white) increased in Dox- condition. (L) FACS analysis of PDGFRα expression on chimeric aggregates. Blue, PKA-ESCs. Green, Control-ESCs. (M) Percentage of differentiating chimeric aggregates expressing PDGFRα. Data represent means ± S.D.

**Figure S2.**
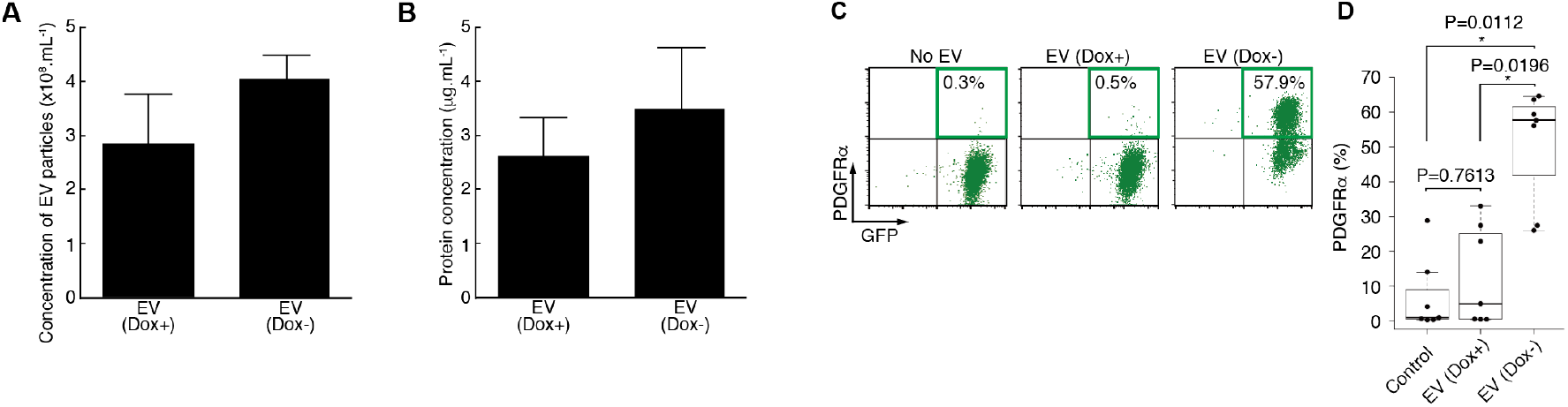
Concentration of EV particles and EV proteins secreted from PKA-ESCs, and mesodermal differentiation of EV-treated Control-ESCs. (A and B) Concentration of EV particles (A) and proteins contained in EVs (B) in conditioned medium. (C) FACS analysis of PDGFRα expression on Control-ESCs at differentiation day 3.5 and treated with EVs from the conditioned medium (Dox+ or Dox-) of PKA-ESCs. (D) Percentage of PDGFRα^+^ cells. Data were analyzed using the Kruskal-Wallis test followed by the Steel-Dwass test. Data represent means ± S.D.

**Figure S3.**
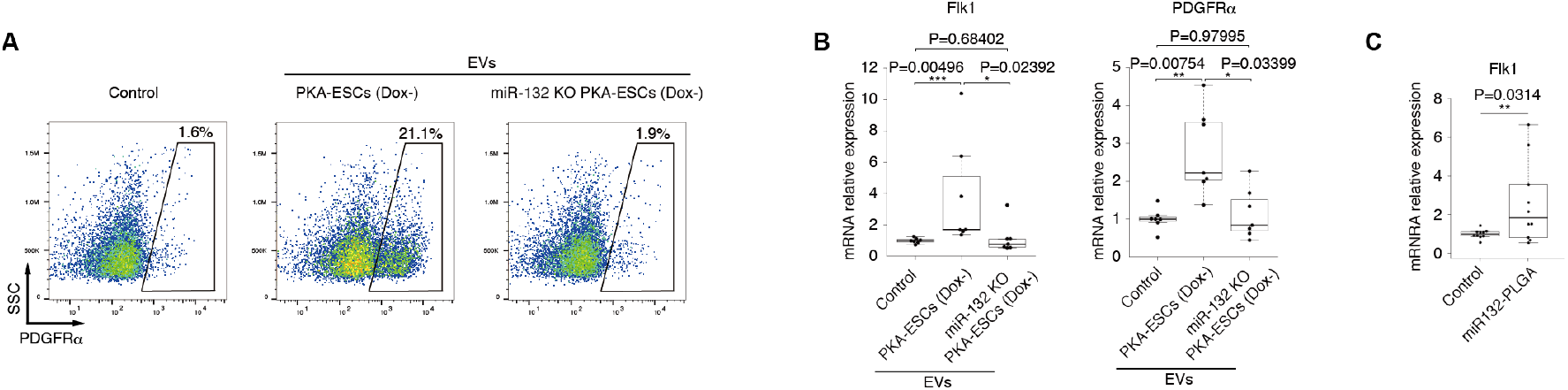
Knockout of miR-132 and mimicking of EV function by miR-132-PLGA. (A) FACS analysis of PDGFRα expression on Control-ESCs at differentiation day 2.5 and treated with EVs from conditioned medium (Dox-) of PKA-ESCs, miR-132-KOPKA-ESCs or control (without EVs). (B) RNA expression levels of Flk1 and PDGFRα in Control-ESCs at day 3.5 treated with EVs from the conditioned medium (Dox-) of PKA-ESCs, miR-132-KO PKA-ESCs or control (without EVs). The values were normalized to control. Data were analyzed using the Kruskal-Wallis test followed by the Steel-Dwass test. (C) mRNA expression levels of Flk1 in Control-ESCs at day 3.5 treated with or without miR-132-PLGA. The values were normalized to control. Data were analyzed using the Mann-Whitney U test.

## References

[1] Arnold, S.J. and Robertson, E.J. (2009). Making a commitment: cell lineage allocation and axis patterning in the early mouse embryo. Nat. Rev. Mol. Cell Biol. 10, 91–103.

[2] Evans, M.J. and Kaufmant, M.H. (1981). Establishment in culture of pluripotential cells from mouse embryos. Nature 292, 154–156.

[3] Martin, G.R. (1981). Isolation of a pluripotent cell line from early mouse embryos cultured in medium conditioned by teratocarcinoma stem cells. Proc. Natl. Acad. Sci. USA 78, 7634–7638.

[4] Thomson, J.A., Itskovits-Eldor, J., Shapiro, S.S., Waknitz, M.A., Swiergial, J.J., Marshall, V.S. and Jones, J.M. (1998). Embryonic Stem Cell Lines Derived from Human Blastocysts. Science 282, 1145–1148.

[5] Yamamizu, K., Kawasaki, K., Katayama, S., Watabe, T. and Yamashita, J.K. (2009). Enhancement of vascular progenitor potential by protein kinase A through dual induction of Flk-1 and Neuropilin-1. Blood 114, 3707–16.

[6] Minakawa, T., Kanki, Y., Nakamura, K. and Yamashita, J.K. (2020). Protein kinase A accelerates the rate of early stage differentiation of pluripotent stem cells. Biochem. Biophys. Res. Commun. 524, 57–63.

[7] Yáñez-Mó, M., Sijander, P. R-M., Andreu, Z., Zavec, A.B., Borràs, F.E., Buzas, E.I., Buzas, K., Casal, E., Cappello, F., Carvalho, J., et al. (2015). Biological properties of extracellular vesicles and their physiological functions. J. Extracell. Vesicles 4, 27066.

[8] Era, T. and Witte, O. (2000). Regulated expression of P210 Bcr-Abl during embryonic stem cell differentiation stimulates multipotential progenitor expansion and myeloid cell fate. Proc. Natl. Acad. Sci. USA 97, 1737–42.

[9] Woltjen, K., Michael, I.P., Mohseni, P., Desai, R., Mileikovsky, M., Hämäläinen, R., Cowling, R., Wang, W., Liu, P., Gertsenstein, M., et al. (2009). piggyBac transposition reprograms fibroblasts to induced pluripotent stem cells. Nature 458, 766–70.

[10] Kataoka, H., Takaura, N., Nishikawa, S., Tsuchida, K., Kodama, H., Kunisada, T., Risau, W., Kita, T. and Nishikawa, S.I. (1997). Expressions of PDGF receptor alpha, c-Kit and Flk1 genes clustering in mouse chromosome 5 define distinct subsets of nascent mesodermal cells. Dev. Growth. Differ. 39, 729–40.

[11] Bedzhov, I., Leung, C., Bialecka, M. and Zernicka-Goetz, M. (2014). In vitro culture of mouse blastocysts beyond the implantation stages. Nat. Protoc., 9, 2732–9.

[12] Kawashima, Y., Yamamoto, H., Takeuchi, H., Hino, T., Niwa, T. (1998). Properties of a peptide containing DL-lactide/glycolide copolymer nanospheres prepared by novel emulsion solvent diffusion methods. Eur. J. Pharm. Biopharm. 45, 41–48.

[13] Yamashita, J., Itoh, H., Hirashima, M., Ogawa, M., Nishikawa, S., Yurugi, T., Naito, M., Nakao, K. and Nishikawa, S. (2000). Flk1-positive cells derived from embryonic stem cells serve as vascular progenitors. Nature 408, 92–96.

[14] Meldolesi, J. (2018). Exosomes and ectosomes in intercellular communication. Curr. Biol. 28, R435–R444.

[15] Dang, L.T.H., Lawson, N.D. and Fish, J.E. (2013). MicroRNA control of vascular endothelial growth factor signaling output during vascular development. Arterioscler. Thromb. Vasc. Biol. 33, 193–200.

[16] Zhang, Y., Liu, Y., Liu, H. and Tang, W. H. (2019). Exosomes: biogenesis, biologic function and clinical potential. Cell. Biosci. 9: 19.

[17] Lei, Z., Mil, A.V., Brandt, M.M., Grundmann, S., Hoefer, I., Smits, M., Azzouzi, H.E., Fukao, T., Cheng, C., Doevendans, P.A., et al. (2015). MicroRNA-132/212 family enhances arteriogenesis after hindlimb ischaemia through modulation of the Ras-MAPK pathway. J. Cell. Mol. Med. 19, 1994–2005.

[18] Gu, X., Su, X., Jia, C., Lin, L., Liu, S., Zhang, P., Wang, X. and Jiang, X. (2018). Sprouty1 regulates neuritogenesis and survival of cortical neurons. J. CELL. PHYSIOL. 234, 12847–12864.

[19] Ding, Q., Gros, R., Gray, I.D., Taussig, R., Ferguson, S.S.G. and Feldman, R.D. (2004). Raf kinase activation of adenylyl cyclases: isoform-selective regulation. Mol. Pharmacol. 66, 921–928.

[20] Sassone-corsi, P. (2012).The cyclic AMP pathway. Cold Spring Harb. Perspect Biol. 4, a011148.

[21] Wang, H., Xu, J., Lazarovici, P., Quirion, R. and Zheng, W. (2018). cAMP response element-binding protein (CREB): A possible signaling molecule link in the pathophysiology of schizophrenia. Front Mol. Neurosci. 11, 255.

[22] Mebratu, Y. and Tesfaigzi, Y. How (2009). ERK1/2 activation controls cell proliferation and cell death is subcellular localization the answer? Cell Cycle 8, 1168–75.

[23] Rasmussen, T.L., Shi, X., Wallis, A., Kweon, J., Zirbes, K.M., Koyano-Nakagawa, N. and Garry, D.J. (2012).VEGF / Flk1 signaling cascade transactivates Etv2 gene expression. PLoS One 7, e50103.

[24] Kataoka, H. Hayashi, M., Nakagawa, R., Tanaka, Y., Izumi, N., Nishikawa, S., Jakt, M.L., Tarui, H. and Nishikawa, S.I. (2011). Etv2 / ER71 induces vascular mesoderm from Flk1^+^PDGFRα^+^ primitive mesoderm. Blood 118, 6975–6987.

[25] Lin, L., Chiu, S., Wu, M., Chen, P. and Yen, J. (2012). Luteolin induces microRNA-132 expression and modulates neurite outgrowth in PC12 cells. PLoS One 7, e43304.

[26] Crewe, C., Joffin, N., Rutkowski, J.M. Kim, M., Zhang, F., Towler, Gordillo, R. and Scherer, P. (2018). An Endothelial-to-Adipocyte Extracellular Vesicle Axis Governed by Metabolic State. Cell 175, 695–708.

[27] van Niel, G., D’Angelo, G. and Raposo, G. (2018). Shedding light on the cell biology of extracellular vesicles. Nat. Rev. Mol. Cell Biol. 19, 213–228.

[28] Hessvik, N.P. and Llorente, A. (2017). Current knowledge on exosome biogenesis and release. Cell Mol. Life Sci. 75, 193–208.

[29] Riva, P., Battaglia, C. and Venturin, M. (2019). Emerging Role of Genetic Alterations Affecting Exosome Biology in Neurodegenerative Diseases. Int. J. Mol. Sci. 20, 4113.

[30] Nabhan, J.F., Hu, R., Oh, R.S., Cohen, S.N. and Lu, Q. (2012). Formation and release of arrestin domain-containing protein 1-mediated microvesicles (ARMMs) at plasma membrane by recruitment of TSG101 protein. Proc. Natl. Acad. Sci. U. S. A. 109, 4146–4151.

[31] Wagner, K.U., Krempler, A., Qi, Y., Park, K., Henry, M.D., Triplett, A.A., Riedlinger, G., Rucker III, E.B. and Hennighausen, L. (2003). Tsg101 is essential for cell growth, proliferation, and cell survival of embryonic and adult tissues. Mol. Cell. Biol. 23, 150–162.

[32] Stoffel, W., Jenke, B., Blöck, B., Zumbansen, M. and Koebke, J. (2005). Neutral sphingomyelinase 2 (smpd3) in the control of postnatal growth and development. Proc. Natl. Acad. Sci. U. S. A. 102, 4554–4559.

